# eggNOG-mapper v2: Functional Annotation, Orthology Assignments, and Domain Prediction at the Metagenomic Scale

**DOI:** 10.1101/2021.06.03.446934

**Authors:** Carlos P. Cantalapiedra, Ana Hernández-Plaza, Ivica Letunic, Peer Bork, Jaime Huerta-Cepas

## Abstract

Even though automated functional annotation of genes represents a fundamental step in most genomic and metagenomic workflows, it remains challenging at large scales. Here, we describe a major upgrade to eggNOG-mapper, a tool for functional annotation based on precomputed orthology assignments, now optimized for vast (meta)genomic data sets. Improvements in version 2 include a full update of both the genomes and functional databases to those from eggNOG v5, as well as several efficiency enhancements and new features. Most notably, eggNOG-mapper v2 now allows: (i) de novo gene prediction from raw contigs, (ii) built-in pairwise orthology prediction, (iii) fast protein domain discovery, and (iv) automated GFF decoration. eggNOG-mapper v2 is available as a standalone tool or as an online service at http://eggnog-mapper.embl.de.

## Main

Inference of gene function via fine-grained orthology, rather than by mere homology detection, is generally considered the most reliable approach for transferring functional information between molecular sequences (Gabaldón and Koonin 2013; Glover et al. 2019). However, since delineating orthology is highly demanding (both computationally and algorithmically), most automated methods rely on homology-based annotations pulled from relatively small reference-species genomic databases (Götz et al. 2008; Jones et al. 2014; Seemann 2014; Shaffer et al. 2020). The eggNOG-mapper method was originally proven to provide more accurate predictions than homology-based approaches (Huerta-Cepas et al. 2017), while preserving computational performance at the genomic and metagenomic scale. This is achieved by inferring fast orthology calls based on precomputed phylogenies, covering thousands of organisms, and a comprehensive database of functional annotations for reference orthologous sequences. Here we present eggNOG-mapper v2, a major upgrade featuring improvements in annotation coverage, overall performance, and program capabilities (**Figure 1A**).

**Figure 1.**
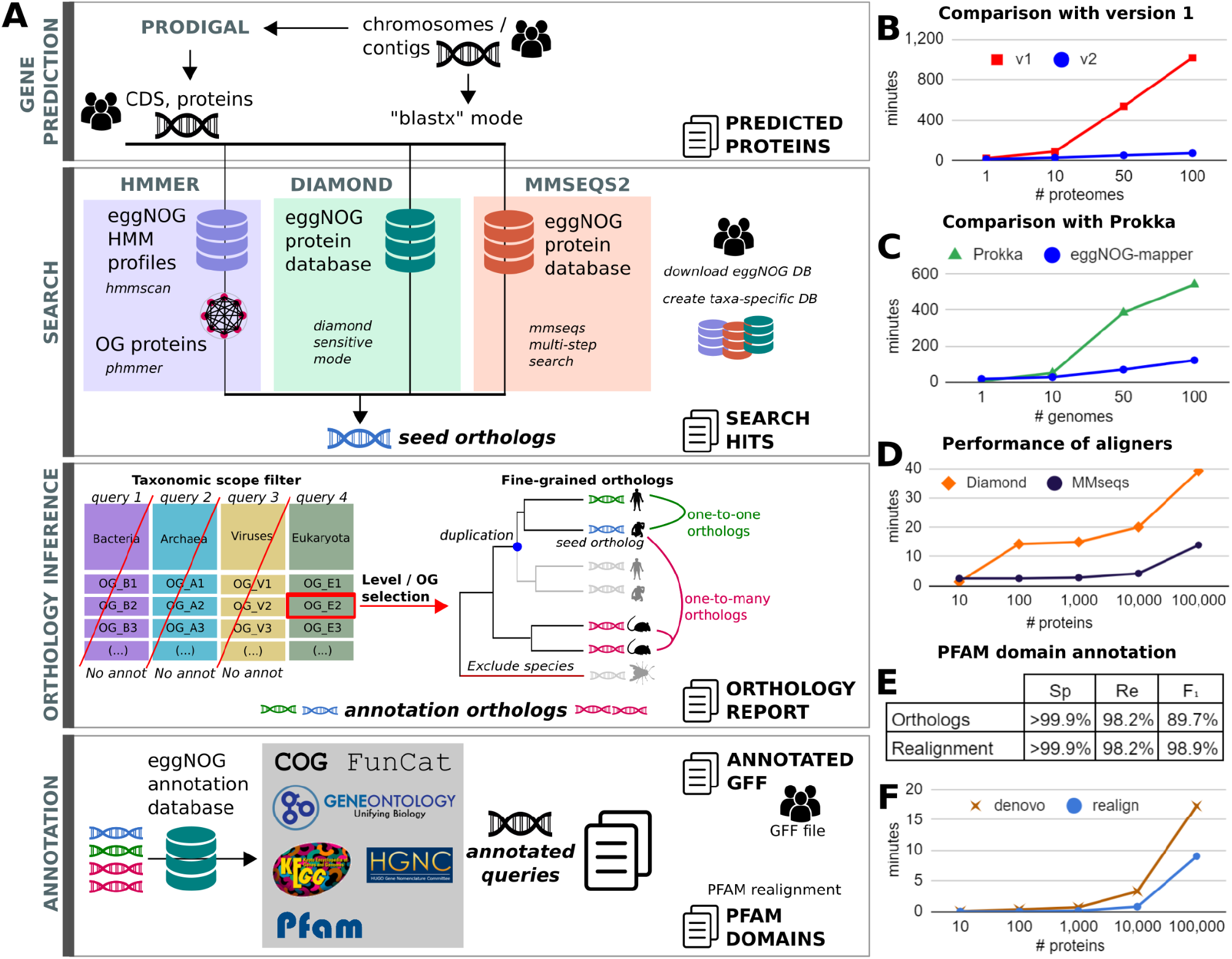
Workflow and performance of eggNOG-mapper v2. **A**: eggNOG-mapper v2 workflow and new features. The gene prediction stage uses Prodigal to perform protein prediction from assembled contigs. During the search stage, HMMER3, Diamond, or MMseqs2 can be used to align the input proteins to eggNOG v5. During the orthology inference stage, a report of orthologs is generated based on the desired taxonomic scope. Finally, protein annotations and domains are transferred from orthologs to the queries and reported as tabular and GFF files. **B**: average minutes to annotate input proteomes. EggNOG-mapper v2 (blue) against eggNOG-mapper v1 (red). **C**: average minutes to annotate input genomes. EggNOG-mapper v2 (blue) against Prokka (green). **D**: average minutes to annotate input proteins. MMseqs2 (-s 3,5,7; black) against Diamond (sensitive mode; orange). **E**: Specificity (Sp), recall (Re), and F_1_ score, of PFAM domain annotation either from direct transference from orthologs, or after realignment. Full de novo realignment results were used as reference. **F**: average minutes for PFAM domain annotation, using either PFAM full de novo (brown) or realign to orthologs domains (blue) modes. **Benchmark setup:** tests in B and C were done on 20 sets of 1–100 random proteomes (B) or genomes (C) from (Almeida et al. 2021), and executed using 10 CPUs and 80GB of RAM. Tests in D, E, and F were done on 25 random sets of 10–100,000 proteins from Progenomes v2 (Mende et al. 2020), using 30 CPUs and 240GB of RAM.

### Coverage and performance improvements

The underlying genome database has been updated to be in sync with eggNOG v5 (Huerta-Cepas et al. 2019), spanning 4.4 million Orthologous Groups (OGs) and more than twice the number of organisms than in the previous version. This improvement increases annotation coverage and phylogenetic resolution, particularly noticeable when analyzing large metagenomic datasets. For instance, the re-annotation of 1.75 million proteins randomly subsampled from a human-gut metagenomic gene catalog (Almeida et al. 2021) yielded a 3.23% increase in annotation coverage (56,569 newly annotated proteins), compared to eggNOG-mapper version 1. Phylogenetic resolution was also improved, obtaining significantly better alignment scores for the query sequences than previous versions (Wilcoxon test W = 1.2E+12, p-value < 2.2E−16). Moreover, although the underlying databases have doubled in size, eggNOG-mapper v2 improves the annotation rate (annotated queries per second) by 16% on average, compared to previous versions. Most important changes regarding computational enhancements relate to database optimizations, allowing for faster queries and parallelization, and a new memory-based mode that significantly reduces the impact of slow I/O disk operations. Taken together, these changes improve annotation rates by 608% on average, with respect to eggNOG-mapper v1 (**Figure 1B**). Compared to Prokka (Seemann 2014), one of the fastest annotation tools available for prokaryotic genomes according to recent benchmarks (Shaffer et al. 2020), eggNOG-mapper runs faster, especially on large metagenomic datasets (**Figure 1C**).

### ORF prediction

Another major capability added to the new eggNOG-mapper workflow is predicting ORFs directly from assembled contigs (“gene prediction” block, **Figure 1A**). ORF detection, only available for prokaryotic assemblies, is performed using Prodigal (Hyatt et al. 2010), which provides the protein sequences to be used by eggNOG-mapper for functional annotation. Prodigal modes (“normal”, “anonymous”, and “training”) as well as custom translation tables can be further chosen by the user.

### Sequence Mapping modes

Additionally, we have broadened the options for the initial sequence-mapping step carried out by eggNOG-mapper (“search” block, **Figure 1A**). Now, Diamond, MMseqs2, and HMMER3 (Mistry et al. 2013) modes are available, each recommended for different use cases. The default Diamond mode provides the best balance between speed and memory consumption. The MMseqs2 mode provides faster results than Diamond (**Figure 1D**) for comparable sensitivity, but requires larger amounts of memory. When input data are nucleotide sequences, a direct translation is done assuming they represent coding sequences starting in an open reading frame. Alternatively, both Diamond and MMseqs2 can be executed in blastx-like mode (for instance, when using sequencing reads as input data). The HMMER3 mode is significantly slower than the other two and requires heavy databases to be downloaded. However, HMM-based searches might increase the annotation coverage for organisms or protein families with distant homology relationships to the eggNOG v5 OGs.

### Adjusting Taxonomic Scopes

Another new feature now available with eggNOG-mapper v2 is the possibility of creating custom annotation databases constrained to specific taxonomic groups. For instance, users could easily create databases spanning only their domain or phylum of interest, therefore reducing computational times of subsequent annotation jobs. Moreover, the new version provides enhanced options to control the taxonomic scope (“orthology inference” block, **Figure 1A**) used for transferring functional annotations, which can be adjusted from automatic mode (recommended for mixed metagenomic datasets) to lineage specific scopes (preventing transferring functional terms from orthologs of unwanted lineages).

### Orthology reports

Taking advantage of the rapid orthology assignments performed by eggNOG-mapper, it is now possible to report pairwise orthology relationships for each query against any of the genomes covered by eggNOG v5 (“orthology inference” block, **Figure 1A**). While this feature is not intended to substitute more precise orthology prediction methods, it provides a very quick and simple “first-pass” approach to obtain pairwise relationships between query sequences and all eggNOG v5 organisms. Orthology reports can be further adjusted by specifying the target taxa and the type of orthologs to be reported (i.e., one-to-one, many-to-many).

### Annotation sources

In order to provide an integrated report of functional annotations per query, eggNOG-mapper v2 offers new annotation sources and improved reports (“annotation” block, **Figure 1A**). The functional annotation sources, which provide different levels of coverage (Supplementary Figure 1), are: predicted protein name; KEGG pathways, modules, and orthologs (Kanehisa et al. 2016); Gene Ontology labels (The Gene Ontology Consortium 2018); EC numbers, BiGG reactions (Norsigian et al. 2019); CAZy terms (Lombard et al. 2014); COG functional categories (Tatusov et al. 2000); eggNOG OGs; and free text descriptions at all taxonomic levels. Reports are generated in tab delimited and/or XLSX file formats. Moreover, when ORF prediction mode is enabled, proteins used to annotate are reported in FASTA format, together with a functionally decorated GFF file. Alternatively, eggNOG-mapper annotation reports can be used to decorate any custom GFF file.

### Protein domain annotations

Along with the functional terms annotated per query, this new version of eggNOG-mapper provides PFAM (Mistry et al. 2020) and SMART (Letunic et al. 2021) protein domain predictions. PFAM domain annotations are by default transferred from the inferred orthologs, which has very little impact on computational cost, but incurs into a small proportion of false positive and negative predictions (F_1_ score 89.7%, **Figure 1E**). Optionally, *de novo* PFAM domain annotation is also available at large scales, both as a refinement phase for the orthology-based predictions (thus keeping computational cost very low, while eliminating the risk of false positives; F_1_ score 98.9%, **Figure 1E**), or by full computation (obtaining native results independent from orthology predictions). When using the *de novo* approach, HMMER3 searches are executed using in-memory mode for higher efficiency. Moreover, GA-based thresholds and PFAM clan disambiguation are automatically applied. Performance comparisons between the different modes are shown in **Figure 1F**.

## Conclusions

Overall, eggNOG-mapper v2 provides a more efficient, versatile, and scalable automated functional annotation workflow than its predecessor. Standalone versions, extended documentation and examples of usage for very large annotation projects (e.g., metagenomic catalogs) are provided at https://github.com/eggnogdb/eggnog-mapper. An online service offering annotation jobs, including thousands of sequences, is available at http://eggnog-mapper.embl.de.

---

**Supplementary Figure 1.**
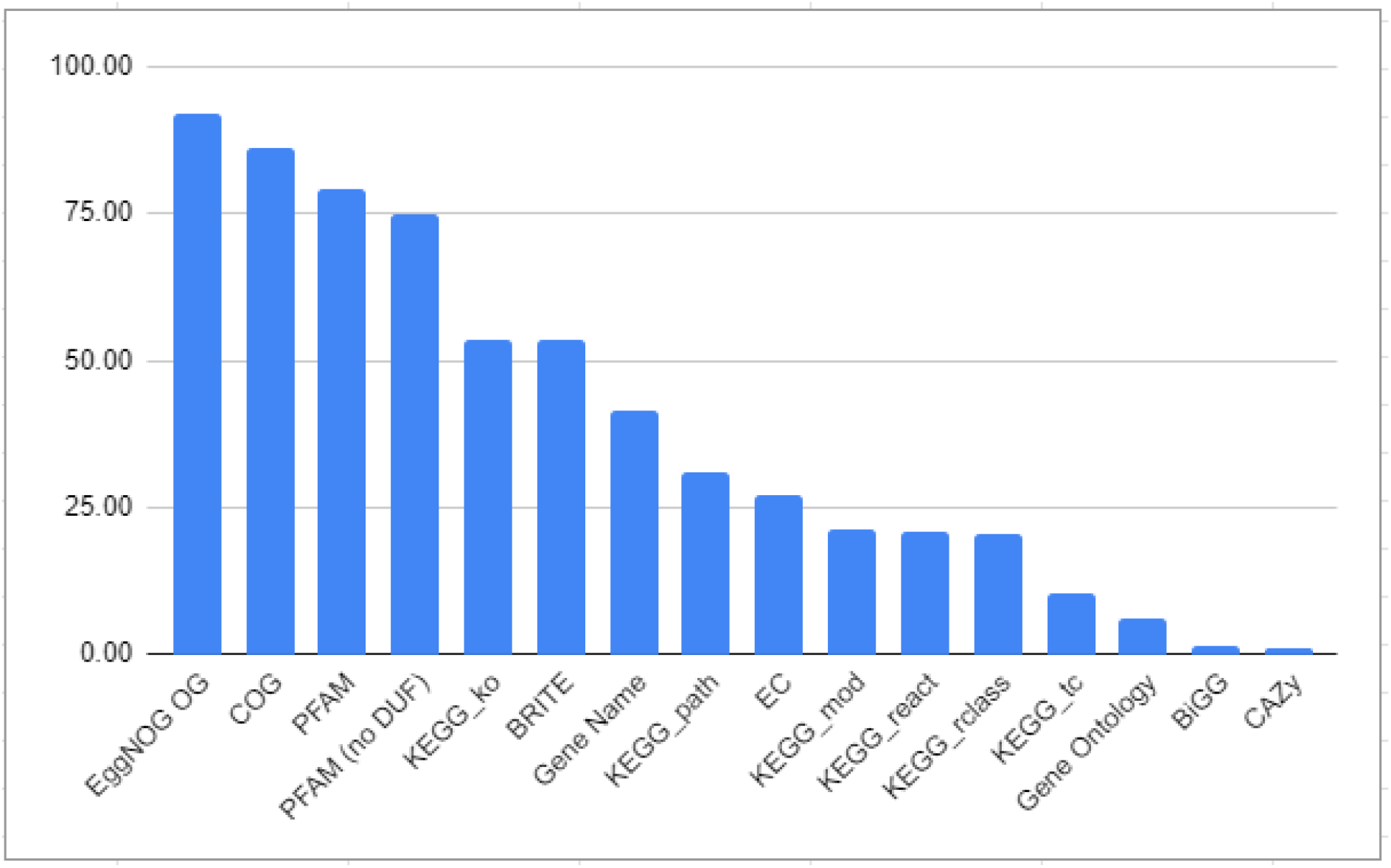
Functional annotation sources included in eggNOG-mapper v2, arranged by the average annotation coverage (% of annotated proteins) in a test with 25 random sets of between 10 and 100,000 proteins from the Progenomes database. PFAM domain annotation obtained from realignment of queries to orthologs domains (‘--pfam_realign realign’ option). The “PFAM (no DUF)” category includes only proteins annotated with at least one non-”DUF” domain.

